# A zebrafish xenotransplant model of anaplastic thyroid cancer to study the tumor microenvironment and innate immune cell interactions *in vivo*

**DOI:** 10.1101/2023.05.29.541816

**Authors:** Cassia Michael, Antonio Di Cristofano, Sofia de Oliveira

**Affiliations:** Department of Developmental and Molecular Biology, Albert Einstein College of Medicine, Bronx, NY, USA; Department of Medicine (Oncology), Albert Einstein College of Medicine, Bronx, NY, USA; Department of Medicine (Hepatology), Albert Einstein College of Medicine, Bronx, NY, USA; Marion Bessin Liver Research Center, Albert Einstein College of Medicine and Montefiore Medical Center, Bronx, NY, USA; Montefiore-Einstein Cancer Research Center, Albert Einstein College of Medicine, Bronx, NY, USA; Cancer Dormancy Tumor Microenvironment Institute, Albert Einstein College of Medicine, Bronx, NY, USA

**Author notes:** **CORRESPONDING AUTHOR:** Sofia de Oliveira.

**Keywords:** Zebrafish, microscopy, tumor microenvironment, neutrophils, xenotransplant

## Abstract

Anaplastic thyroid cancer (ATC) is a rare malignant subtype of thyroid cancer. While ATC is rare it accounts for a disproportionately high number of thyroid cancer-related deaths. Here we developed an ATC xenotransplant model in zebrafish larvae, where we can study tumorigenesis and therapeutic response in vivo. Using both mouse (T4888M) and human (C643) derived fluorescently labeled ATC cell lines we show these cell lines display different engraftment rates, mass volume, proliferation, and angiogenic potential. Next, using a PIP-FUCCI reporter to track proliferation *in-vivo* we observed cells in each phase of the cell cycle. Additionally, we performed long-term non-invasive intravital microscopy over 48 hours to understand cellular dynamics in the tumor microenvironment at the single cell level. Lastly, we tested a well-known mTOR inhibitor to show our model could be used as an effective screening platform for new therapeutic compounds. Altogether, we show that zebrafish xenotransplants make a great model to study thyroid carcinogenesis and the tumor microenvironment, while also being a suitable model to test new therapeutics *in vivo*.

**SUMMARY STATEMENT:** Anaplastic thyroid cancer xenotransplant model in zebrafish larvae to study thyroid cancer tumorigenesis and tumor microenvironment. Using confocal microscopy to understand cell cycle progression, interactions with the innate immune system, and test therapeutic compounds in vivo.

## Introduction

Anaplastic thyroid cancer (ATC) is a rare, undifferentiated, and highly aggressive form of thyroid cancer associated with poor prognosis. Although its incidence is low (i.e., ∼3% of all thyroid cancer cases), it accounts for nearly half of all thyroid cancer-related deaths, as a consequence of its unresponsiveness to current treatment options (1). Since many aspects of the biology of these tumors are still obscure, we decided to develop a novel ATC xenotransplant model in zebrafish larvae that allows to study thyroid cancer biology and therapeutic response in vivo, including evaluation of the tumor architecture, composition and dynamics of the tumor microenvironment, and the visualization of drug effects in vivo.

Zebrafish are a powerful non-mammalian vertebrate model to study many human diseases, including cancer. Xenotransplants in zebrafish larvae are a well-established model for multiple cancers, such as colorectal and breast, that have gained popularity for their fast development, high fecundity, easy genetic manipulation and *in vivo* tractability (2, 3, 4). But perhaps their biggest asset is optical transparency, which allows for long-term non-invasive live imaging of the tumor microenvironment, and the ability to perform robust high throughput drug and genetic screenings (2). Importantly, they offer a shorter experimental timeline and do not have a fully developed adaptive immune system until 2-3 weeks post fertilization, with lymphoid lineage appearing by 10 days post-fertilization (5). This allows for the use of an immune competent model and a unique opportunity to study the interaction between cancer cells and innate immune cells, and further explore mechanisms of innate immune evasion (6). While ATC xenotransplants in zebrafish have been previously reported, a full characterization of the model has never been done (7).

Here, we injected fluorescently labeled mouse and human-derived ATC cell lines into the perivitelline space (PVS) of zebrafish larvae at 2 days post fertilization (dpf) (3) and tracked their engraftment rate, mass volume, proliferation and angiogenic potential. Furthermore, we tested AZD2014, a well-known mTOR inhibitor, to validate its ability to arrest cells in our model. Altogether, our results show that zebrafish larvae xenotransplants are a promising *in vivo* model to investigate ATC tumor microenvironment, tumorigenesis, and explore new therapeutics.

## Results

### ATC cell lines engraft in zebrafish larvae at different rates

Zebrafish are a powerful and dynamic model organism to study cancer biology (6). To test if anaplastic thyroid cancer (ATC) cell lines can engraft 2 days post fertilization (dpf) zebrafish larvae, were injected with mouse N794, T1903, T1860, N2773, N2933, and T4888M and human ACT1, BHT101, ASH3, and C643 ATC cell lines and followed masses up to 4 days post injection (dpi) (Figure 1A). Around 300-500 cancer cells were injected into the perivitelline space (PVS), the pocket between the zebrafish yolk and the outer dermal membrane (Figure 1B), mimicking a subcutaneous injection in mouse model. Next, we explored the engraftment potential of the different ATC cell lines (Figure 1C). At 4dpi, we observed that mouse cell lines have similar engraftment potential, with an engraftment rate ranging between 70%-100 (N794=87%, T1903=69%, T1860=83%, N2773=98%, N2933=60% and T4888M=98%), while human cell lines have engraftment rates between 40-60% (ACT1=57%, BHT101=60%, ASH3=42% and C643=64%) (Figure 1C and D). Histopathological analysis of the zebrafish tumor masses derived from the T4888M cell line shows that they have similar morphological features as observed previously in mice (Figure 1E-G). In addition, we observed two major characteristics of ATC thyroid cancer in both T4888M and C643 zebrafish masses, which are spindle cell/nuclear spindle formation and pleomorphic cells (11). Spindle cell and nuclear spindle formation can be identified by their elongated and thinner morphology (Figure 1E’-G’, 1L’), while pleomorphic cells were identified by their abnormal shape and variation amongst cells (Figure 1E’-G’, 1J’-L’). These findings suggest that both human and mouse ATC cell lines can engraft in zebrafish larvae and that masses preserve major ATC histological features.

**Figure 1.**
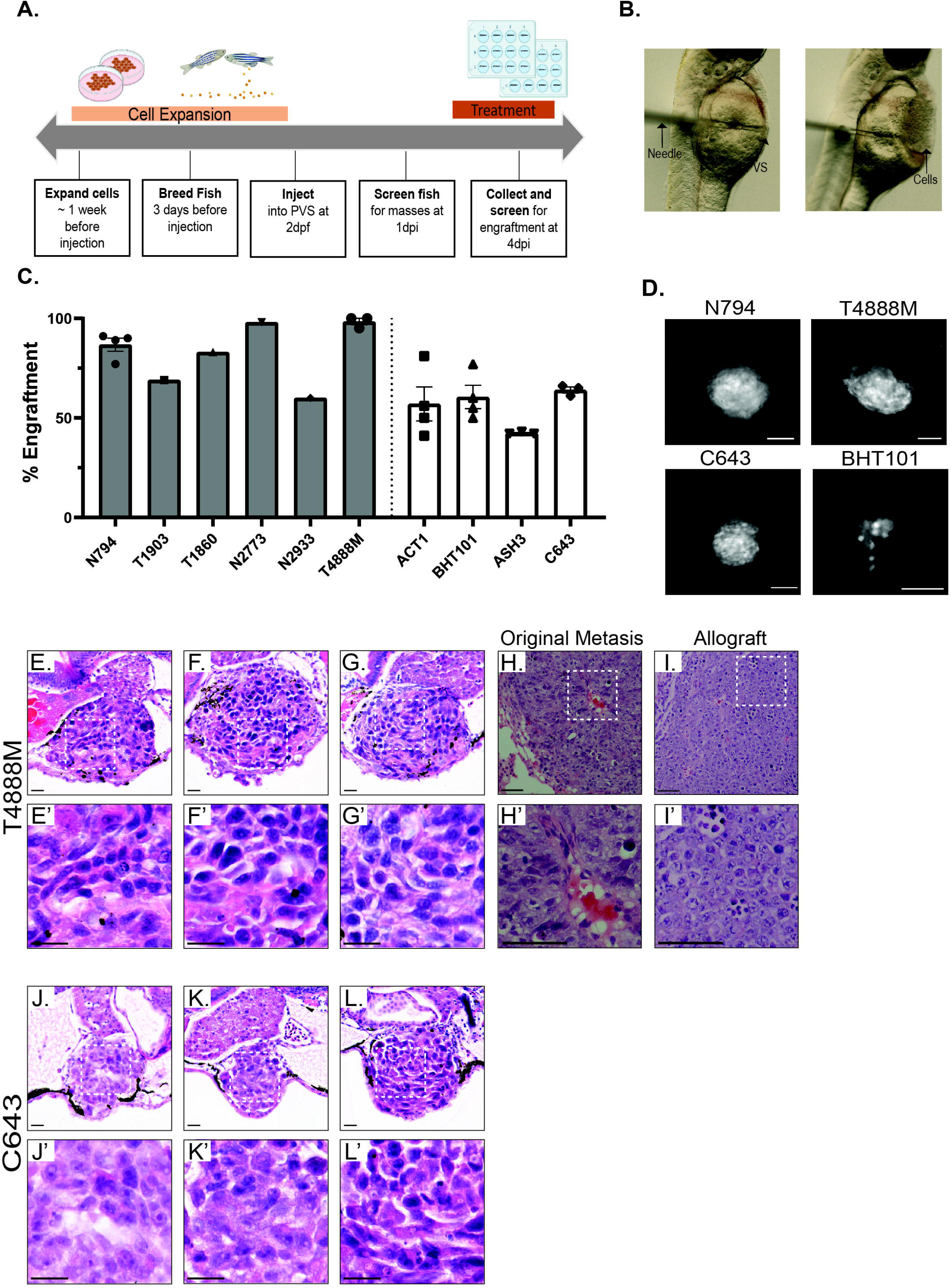
ATC cell lines engraft in zebrafish larvae at different rates. (A) Experimental timeline for thyroid xenotransplant model. (B) Representative images acquired using the stereomicroscope at 2 dpf to show proper site of injection. (C) Graph showing percent engraftment for ATC mouse (N794, T1903, T1860, N2773, N2933, T4888M) and human (ACT1, BHT101, ASH3, C643) cell lines. (D) Representative images acquired using the fluorescent microscope at 4 dpi in mouse and human ATC cell lines. Scale bar= 100 μm (E-G & J-L) Representative images of T4888 and C643 ATC cell lines in zebrafish. Scale bar= 20 μm. (E’-G’ & J’-L’) Higher magnification showcasing important histopathological features maintained in T4888 and C643 ATC zebrafish xenotransplants. Scale bar= 20 μm. (H & H’) Histopathological analysis from original lung metastasis of T4888 ATC cells in a murine mouse model ((Pten, P53)/-/-). Scale bar= 50 μm. (I & I’) Histopathological analysis from an allograft in a murine mouse model. Scale bar= 50 μm. Data is from at least two independent experimental replicates. Bar plots show mean ± SEM, each dot represents one experiment.

### T4888M and C643 ATC cell lines display different growth and proliferation rates

For the remainder of the experiments, we utilized the two cell lines with higher engraftment rates, the mouse line T4888M and the human line C643. To further characterize the ATC xenografts, we next tested if both human and mouse lines could grow and proliferate in the zebrafish. Thus, we performed non-invasive intravital confocal microscopy of xenografts at 1 and 4dpi and after IMARIS imaging analysis we observed that the T4888M cell line mass volume significantly increased over the 4 days while a reduction was observed for the C643 cell line (Figure 2A,2B). Additionally, we assessed the proliferation of larvae xenografts for both cell lines by EDU (5-ethynyl-2’deoxyuridine) incorporation over 4 hours. Not surprisingly, we observed a higher density and number of EDU positive cells in the T4888M xenograft masses in comparison to the C643 (Figure 2E,2F). Overall, our results show that both mouse and human lines can proliferate in the zebrafish larvae, with mouse ATC line demonstrating higher growth and proliferative rates, while in human cells the proliferative rate is not sufficient to overcome cell loss.

**Figure 2.**
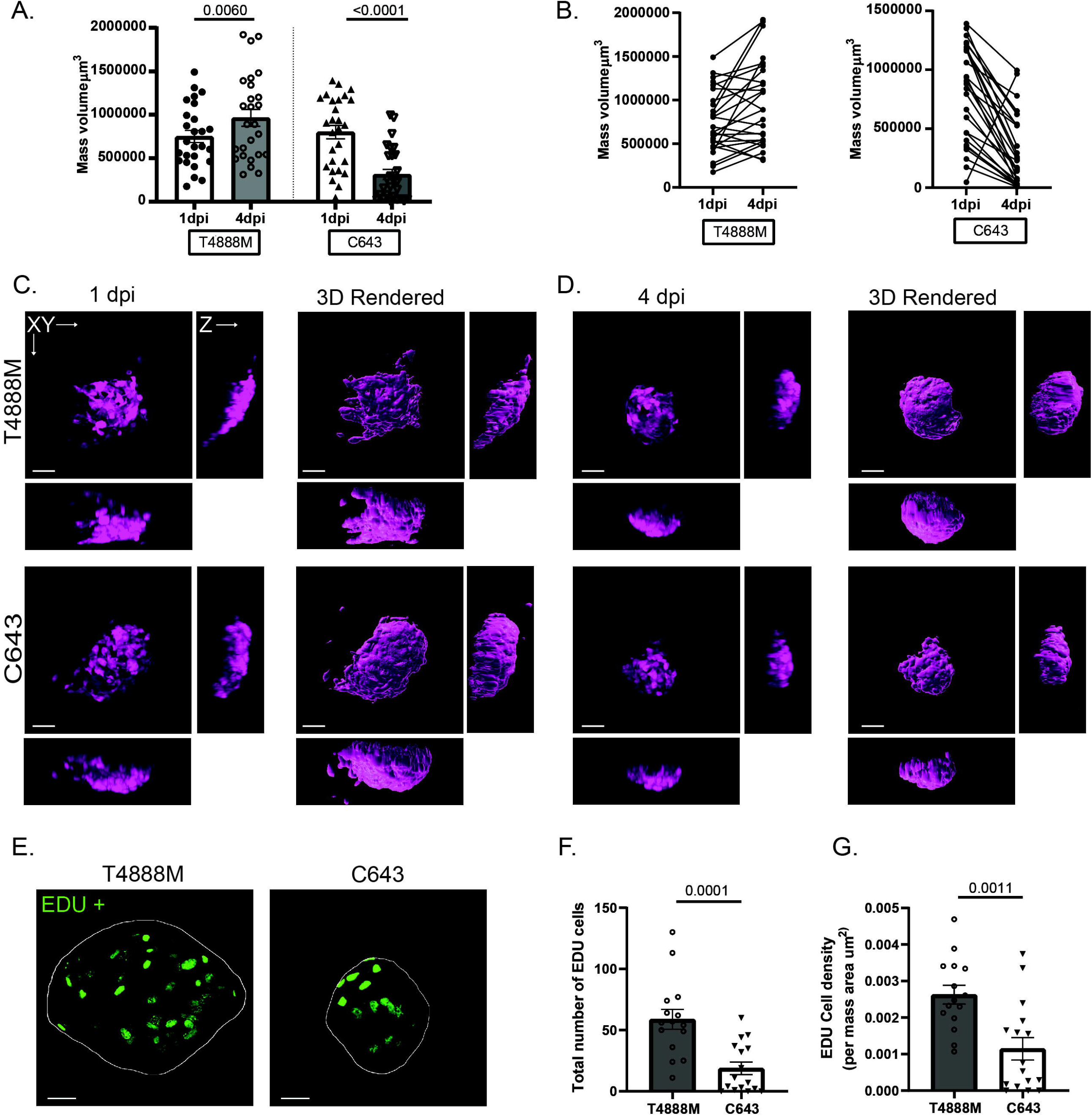
T4888M and C643 ATC cell lines display different growth and proliferation rates. (A) Graph showing mass volume μm^3^ at 1 dpi compared to 4 dpi in ATC cell lines T4888 and C643 (T4888 n=26, C643 n=27). (B) Graph showing mass volume μm^3^ at 1 dpi compared to 4 dpi tracking individual larva in ATC cell lines T4888 and C643. (C-D) Representative 3D images and reconstruction of ATC tumor masses from 1 dpi to 4 dpi in ATC cell lines T4888 and C643. Images were acquired using the 20x objective. Scale bar= 50 μm. (E) Representative 3D images of EDU+ cells inside cancer mass of ATC cell lines T4888 and C643. (F) Quantification of the total number of EDU+ cells in T4888 and C643 ATC masses (T4888 n=15, C643 n=16). Images were acquired using the 40x objective. Scale bar= 30 μm. (G) Quantification of the number of EDU+ cells normalized by the area μm^2^ of the ATC mass. Data is from at least two independent experimental replicates. Bar plots show mean ± SEM, each dot represents one larva, P-values are shown in each graph.

### T4888M ATC xenotransplants display high angiogenic ability

An important feature that supports cancer growth and proliferation is angiogenesis (12). To understand why there are such large differences in mass volume and proliferation between the T4888M and C643 cell lines, we looked at the ability of both cell lines to recruit vessels to the mass using a transgenic zebrafish line, Tg(Fli1:eGFP) that labels vasculature with GFP (13). At 4dpi, we observed that the T4888M cell line has a dramatically higher angiogenic potential, with xenografts displaying highly vascularized masses compared to the C643 cells (Figure 3A-D). The T4888M cell line forms a dense organization of vessels that infiltrate the mass, while the C643 masses recruit almost no vessels to the mass (Movie S1,S2). Correlation analysis shows that increased vessel volume is correlated to increased mass volume for the T4888M xenografts (Figure 3E). However, for the C643 xenografts, vessel volume is not correlated to mass volume, which is consistent with the larger C643 masses we observed that lack vessel infiltration (Figure 3F). Together, these findings suggest that T4888M has an extensive angiogenic potential compared to C643 that may be contributing to the observed higher survival, proliferation and growth rates.

**Figure 3.**
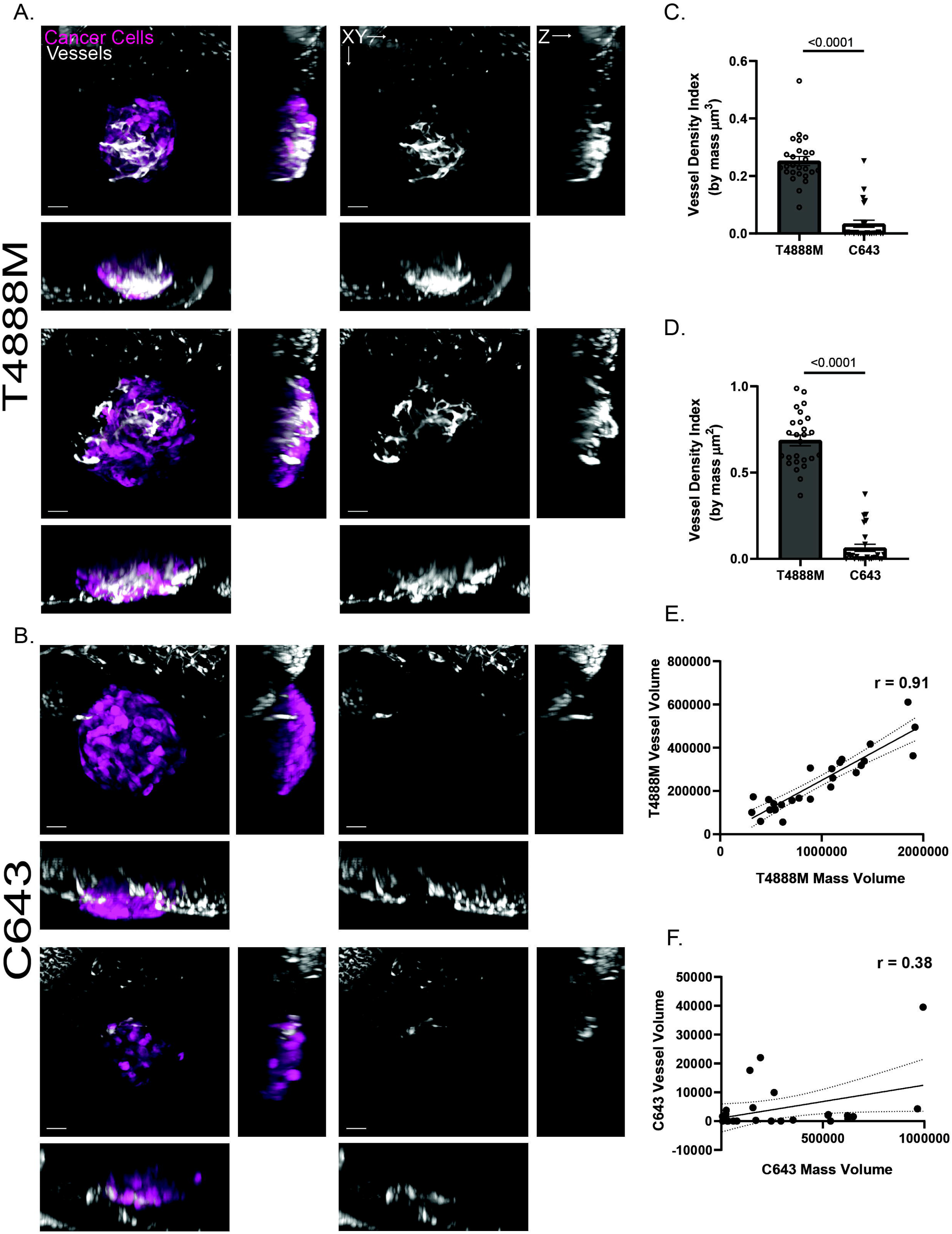
T4888M ATC xenotransplants display high angiogenic ability. (A-B) Representative 3D images of ATC tumor mass and vessel recruitment at 4 dpi Tg(fli:GFP) in ATC cell lines T4888 and C643. Scale bar= 50 μm. (C) Graph showing vessel density in tumor mass normalized by tumor volume μm^3^ for ATC cell lines T4888 and C643 (T4888 n=26, C643 n=26). (D) Graph showing vessel density in tumor mass normalized by tumor area μm^2^ for ATC cell lines T4888 and C643 (T4888 n=26, C643 n=26). (E) Graph of the correlation between T4888 mass volume and T4888 vessel volume. (F) Graph of the correlation between C643 mass volume and C643 vessel volume (T4888 n=26, C643 n=26). Data is from at least two independent experimental replicates. Bar plots show mean ± SEM, each dot represents one larva, P-values are shown in each graph. Pearson correlation graphs show line of best fit; each dot represents one larva, r-values are shown in each graph.

### In vivo visualization of T4888M cell cycle using the PIP-FUCCI reporter

To further understand the proliferative capacity and cellular dynamics of the T4888M cell line, we incorporated a PIP-FUCCI reporter into the cells. This reporter allowed us to delineate various phases of the cell cycle, optimizing our ability to do long-term in vivo imaging at the single cell level (14). This two-colored reporter expresses GFP when cells are in G1 and as they transition to late S phase it expresses mCherry. When cells are in G2/M both GFP and mCherry are expressed, allowing for colocalization of signal to detect these cells (14). Again, using non-invasive confocal microscopy followed by IMARIS imaging analysis we determined which phase of the cell cycle cells were at 4dpi (Figure 4A). At this timepoint we observe the highest number of cells to be in G1 and the second highest to be in late S-phase (Figure 4B). This is also consistent when the number of cells in each phase is normalized by the total number of cells in each mass (Figure 4C). Next, to investigate the cell division dynamics on T4888 masses, we performed long term time-lapse microscopy for over 48 hours and observed that these masses maintain their ability to proliferate (Figure 4D-E; Movie S3).

**Figure 4.**
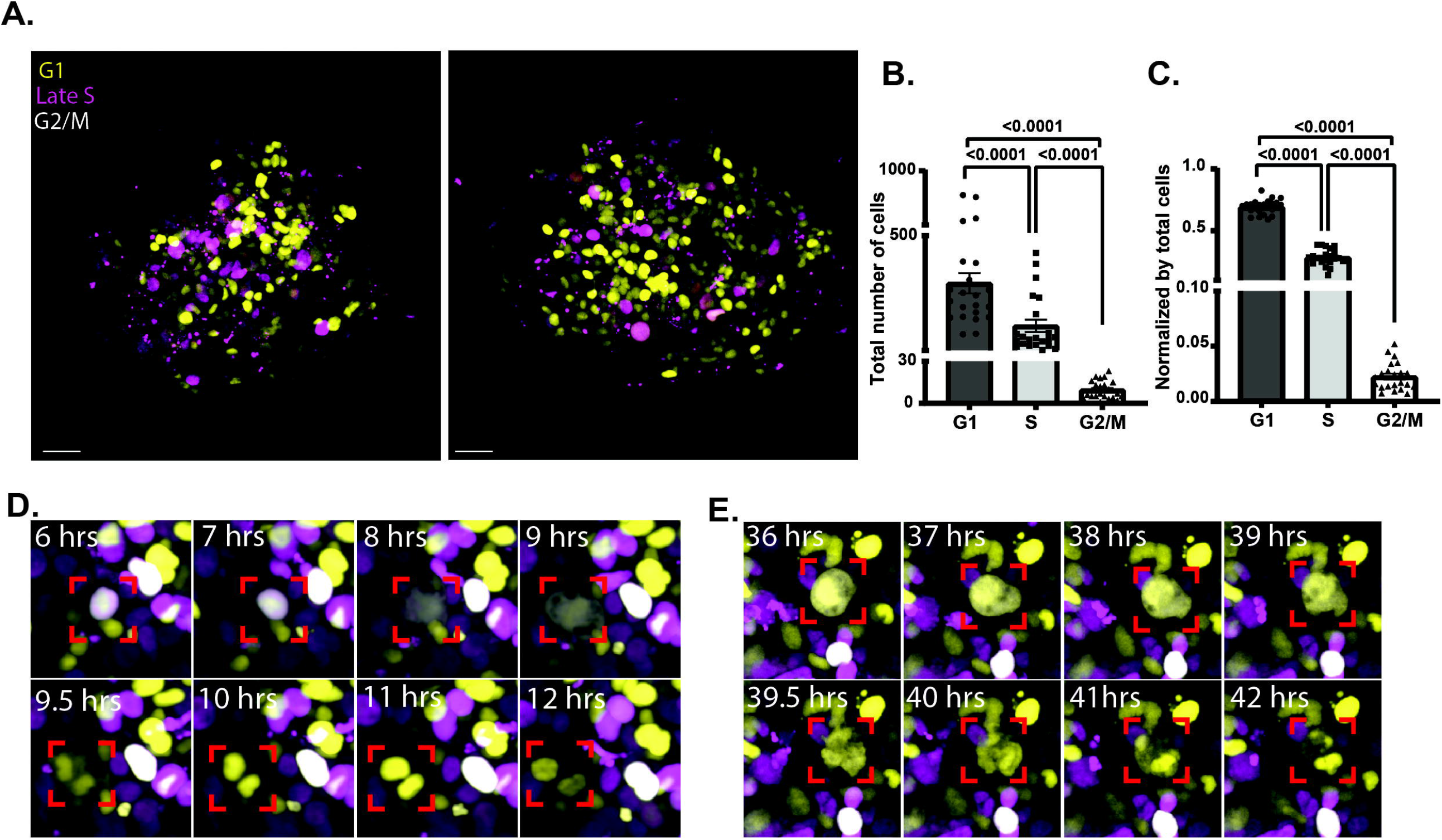
In vivo visualization of T4888M cell cycle using the PIP-FUCCI reporter. (A) Representative 3D images of T4888 ATC cancer cells using the PIP-FUCCI reporter. Scale bar= 30 μm. (B) Graph showing total number of cells at 4 dpi in each cell cycle phase with the PIP-FUCCI reporter (T4888 n=22). (C) Graph showing cell cycle phases with the PIP-FUCCI reporter normalized by the total number of cells in T4888 ATC mass (T4888 n=22). (D-E) Images from 48-hour live imaging to highlight proliferative cells, imaging begins 1dpi and ends at 3dpi. Data is from at least two independent experimental replicates. Bar plots show mean ± SEM, each dot represents one larva, P-values are shown in each graph.

### T4888M ATC cell line responds to cell cycle inhibitors

To understand if our model could be used for preclinical screenings, we performed a proof-of-principle experiment using the well-known mTOR inhibitor, AZD2014. Using the PIP-FUCCI reporter coupled with non-invasive confocal microscopy and image analysis we observed that the inhibitor significantly reduced the total number of cells in each mass (Figure 5A, 5B). When looking more closely at the individual cell phases, we observed that the AZD2014 compound causes a reduction of cells in late S and an increase in cells in G1, when compared to the DMSO control (Figure 5C). As expected, the treatment did not have a significant impact on cells in G2/M phase. Lastly, we looked at neutrophil infiltration into T4888 ATC masses (Figure 5D). We observed neutrophils to infiltrate these masses and after treatment with AZD2014 we observed a reduction in the total number of neutrophils, when compared to the DMSO control (Figure 5E & 5F). This data suggest that our model is a viable and powerful system that can be used to screen compounds that impact ATC tumorigenesis. Additionally, it can be used to study the impact of the innate immune cells on ATC tumorigenesis and the tumor microenvironment.

**Figure 5.**
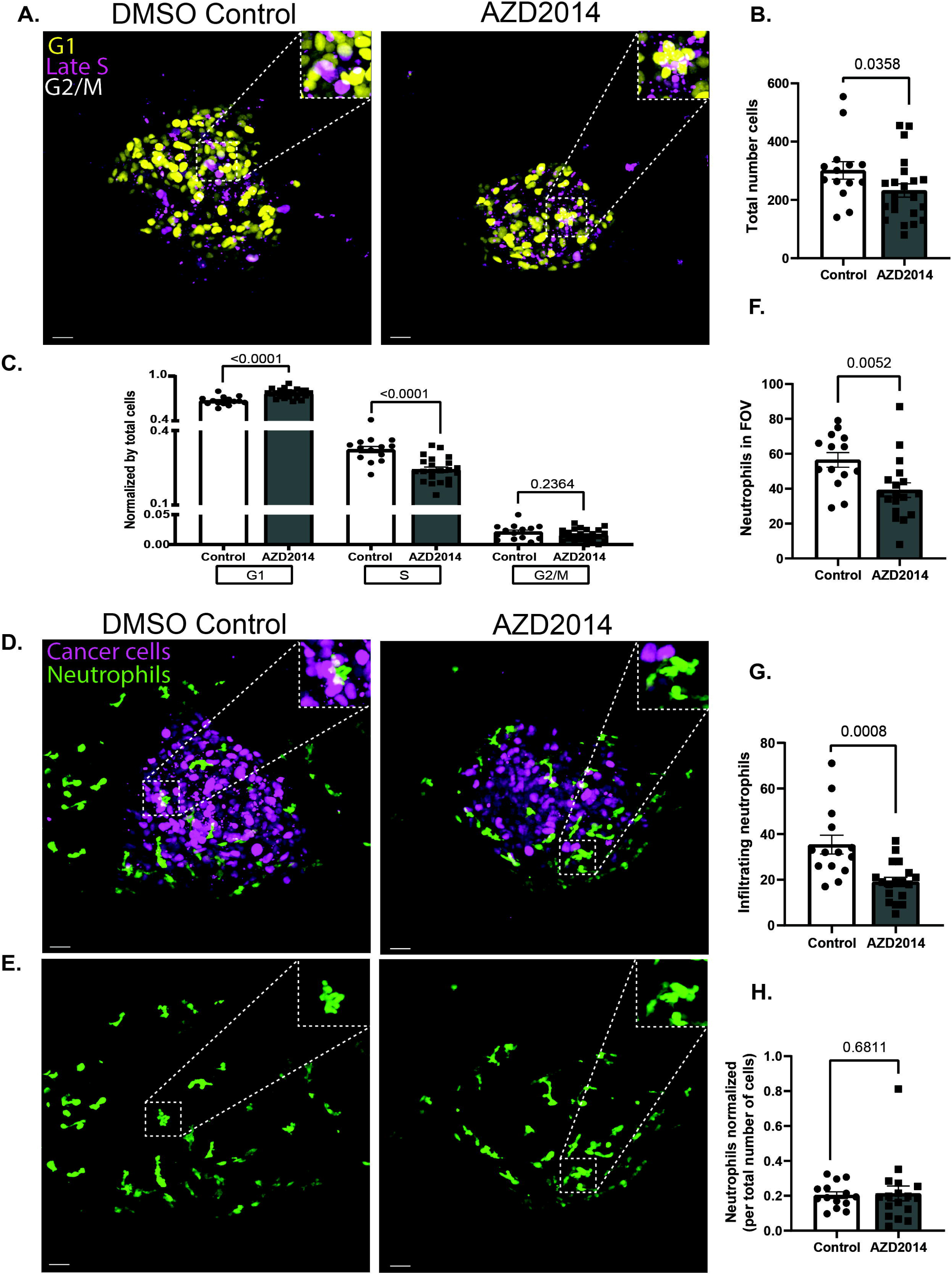
T4888M ATC cell line responds to cell cycle inhibitors. (A-C) Representative 3D images of ATC tumor mass and neutrophil recruitment at 4 dpi Tg(lyzC:BFP) with DMSO control and AZD2014 treatment. Zebrafish larvae were treated with 0/1% DMSO control or 1 μm AZD2014 (Vistusertib). Scale bar= 20 μm. (D) Graph showing cell cycle phases with the PIP-FUCCI reporter normalized by the total number of cells in DMSO control and AZD2014 treatment (DMSO Control n=14, AZD2014 n=22). (E) Graph showing total number of cells in each mass in DMSO control and AZD2014 treatment (DMSO Control n=14, AZD2014 n=22). (F) Graph showing the total number of neutrophils in field of view (FOV) for the DMSO control and AZD2014 (DMSO Control n=14, AZD2014 n=18). (G) Graph showing total number of neutrophils infiltrating the mass in the DMSO control and AZD2014 (DMSO Control n=14, AZD2014 n=18). (H) Graph showing the total number neutrophils normalized by the total number of cells in each larva (DMSO Control n=14, AZD2014 n=18). Data is from at least two independent experimental replicates. Bar plots show mean ± SEM, each dot represents one larva, P-values are shown in each graph.

## Discussion

Anaplastic thyroid cancer (ATC) is the deadliest and most aggressive form of thyroid cancer, with an average survival of 9 months after diagnosis. Although ATC has a low incidence rate, its highly aggressive nature makes it resistant to many already approved therapeutics, generating a need for new and efficient approaches (15). While ATC zebrafish xenotransplant models have been previously reported (7), here we developed and characterized a new in vivo model that allows visualization of cellular dynamics and tumorigenesis non-invasively at the single cell level. Additionally, our model provides an approach for the identification of new small molecules and actionable targets that can effectively inhibit tumor progression.

The ability to perform whole animal non-invasive live imaging, easy genetic manipulation and cell trackability have made zebrafish an extremely powerful system to study cancer biology both using xenotransplants or tumor-induced models (16). As a small vertebrate with more than 80% homology in genes related to human disease, zebrafish can be modulated to further understand cancer related pathways, through transgenesis, gene specific knockouts or by modulating cell responsiveness to the tumor microenvironment (17). Zebrafish larval xenografts have proven to be a powerful model with many advantages over traditional xenograft models. Reduced size scale and time make this a unique system to investigate cancer biology and tumor microenvironment, perform medium-high throughput drug screenings, and explore personalized medicine approaches (2, 17).

Our findings show that zebrafish larvae are a viable model to study thyroid carcinogenesis and the ATC tumor microenvironment *in vivo*. We found that different mouse and human ATC lines successfully engraft, with rates ranging from 40 to 100%. Zebrafish thyroid xenograft models effectively display many hallmark characteristics of tumor progression and ATC histopathology. Spindle cell and nuclear spindle cell formation are the most abundant cytological features seen in ATC cells, identified by their long slender shape that assemble in an almost striated pattern (11). Pleomorphic cells are the second most common cytological feature found in ATC tumors characterized by abnormal cell and nuclear variation (11).Both of these features were observed in xenograft masses, indicating that ATC cells are maintaining their original features. While it’s important for xenotransplants to maintain these characteristics, it is of equal importance for them to sustain proliferative capabilities. One of the most fundamental traits of cancer cells is their ability to proliferate uncontrollably. Here, we show that after cell transplantation both the T4888M and the C643 cancer cells maintain their ability to proliferate in zebrafish larvae in a period over 4 days, however T488M displays a much higher proliferation rate compared to C643. Another hallmark of tumor progression is the ability to recruit vasculature to the mass (12). Tumor masses require recruitment of vasculature to continue growing. Proper vasculature provides oxygen, nutrients, and removes waste from masses. Additionally, it facilitates the dissemination and metastasis of cancer cells to distant sites by access to circulation (18). We observe at 4 dpi that the T4888M xenografts display a high angiogenic rate that positively correlates with mass size, contrary to the C643 line, which shows a low angiogenic potential, recruiting almost no vessels to the masses. This difference in angiogenic potential between these two lines can partially explain the lower proliferative rate and mass growth found in C643. Indeed, the C643 line compared to other ATC cell lines produces lower amounts of IL8, which is a powerful pro-angiogenic factor including in zebrafish model (19, 20). Surprisingly, some C643 xenografts developed big masses with low to no vessel formation within the masses, suggesting that this cell line can grow at some extent without the presence of tumor vasculature, at least in the zebrafish larvae model. It is important to note that some ATC cell lines do not engraft as well as others; which can potentially be explained by mechanism of innate immune cell evasion, as observed for models of colon cancer: future experiments will provide more insight into these differences and mechanism in ATC (3).

Next, we looked at T4888M cell cycle dynamics at the single cell level using a PIP-FUCCI reporter. The PIP-FUCCI reporter delineates each phase of the cell cycle, which gave us a better understanding of T4888M ability to proliferative *in vivo*. Using this tool associated with long term time-lapse confocal microscopy, we observed that T4888M injected cells engage in a dynamic proliferative stage, as shown by EDU incorporation at 4dpi. Next, as a proof of principle, we tested the effect of on proliferation of AZD2014 a dual mTORC1 and mTORC2 inhibitor which is expected to inhibit T4888M cells, which are driven by a mutation in *Pten* that activates the PI3K/AKT/mTOR signaling pathway. We observe, at 3 days post treatment with AZD2014, a reduction in the number of proliferating cells and an increase in the number of cells in G1, suggesting the cells are undergoing arrest. Additionally, we were able to observe that ATC masses are heavily infiltrated with neutrophils, without changes in infiltration density upon treatment with AZD2014. ATC cells may be releasing inflammatory chemokine signals calling neutrophils to the mass, such as CXCL1 or CXCL8 and may be interesting targets for future work (21). Nevertheless, the subtype of neutrophils that are present before and after treatment might be different. Neutrophils play a dual role in cancer depending on the tumor microenvironment. They can be pro-tumor supporting cell cycle progression and metastasis.

But can also acquire a cytotoxic anti-tumor role. Recently, it has been shown neutrophils immune checkpoint (IC) therapy activate neutrophils to clear tumor antigen escape variants and elicit neutrophils to acquire an interferon signature that is essential for successful response to therapy (22). Our zebrafish ATC model will be a valuable tool to help better understand the pathophysiological role these innate immune cells are playing in ATC and if they can be explored as a way to help eradicate ATC.

Overall our work provides a new powerful *in vivo* platform that allows for fast and highly sensitive assays that can be used to better understand ATC biology, tumor microenvironment, and therapeutic efficiency, as well as explore small molecule drug screening (3). In addition, zebrafish xenografts could represent the initial step for preclinical screening before moving to a more expensive model, helping to expedite studies from bench to bedside, and potentially to personalized medicine (5). Taken together, our data suggests that zebrafish are a promising model to study ATC cancer biology in particular when using cell lines derived from relevant genetically engineered mouse models.

## Materials and Methods

### Zebrafish general procedures and lines

All protocols using zebrafish in this study were approved by the Albert Einstein College of Medicine Institutional Animal Care and Use Committees (IACUC). For all experiments, larvae were anesthetized in E3 (embryo medium) without methylene blue containing 0.16 mg/ml tricane (MS22/ethyl3-aminobenzoate - Sigma Aldrich). Transgenic line expressing BFP under the lyzC promoter in the membrane of neutrophils Tg(lyzC:BFP)^zf217Tg^ and transgenic line expressing eGFP under the fli1 promoter of endothelial cells Tg(fli1:EGFP)^y1Tg^ were used in this study (4). The mutant line Casper^Roya9;mitfaw2^ was also used in this study (5).

### ATC Cell line maintenance

All cell lines in this study (Table 1) were maintained at 37°C with 5% CO2 in RPMI1640 (human cells) or DMEM (mouse cells) with 10%FBS. Cell line identity was confirmed by STS profiling as well as by amplifying and sequencing genomic fragments for known mutations (7-8).

**Table 1.**

| **Cell Line** | **Species of origin** | **Mutations** | **Culture Medium** |
| --- | --- | --- | --- |
| N794 | Mouse | *Nras*<sup>Q61R</sup>, *Pten*<sup>-/-</sup>, *Tp53*<sup>-/-</sup> | DMEM |
| T4888M | Mouse | *Pten*<sup>-/-</sup>, *Tp53*<sup>-/-</sup> | DMEM |
| T1903 | Mouse | *Pten*<sup>-/-</sup>, *Tp53*<sup>-/-</sup> | DMEM |
| T1860 | Mouse | *Pten*<sup>-/-</sup>, *Tp53*<sup>-/-</sup> | DMEM |
| N2773 | Mouse | *Nras*<sup>Q61R</sup>, *Pten*<sup>-/-</sup>, *Tp53*<sup>-/-</sup> | DMEM |
| N2933 | Mouse | *PIK3CA*<sup>H1047R</sup>, *Pten*<sup>-/-</sup>, *Tp53*<sup>-/-</sup> | DMEM |
| BHT101 | Human | *BRAF*<sup>V600E</sup>, *TP53*<sup>I251T</sup> | RPMI |
| CAL62 | Human | *KRAS*<sup>G12R</sup>, *TP53*<sup>A161D</sup> | DMEM |
| C643 | Human | *HRAS*<sup>G13R</sup>, *TP53*<sup>R248Q</sup> | RPMI |
| OCUT2 | Human | *PIK3CA*<sup>H1047R</sup>, *BRAF*<sup>V600E</sup> | RPMI |
| ACT1 | Human | *NRAS*<sup>Q61K</sup>, *TP53*<sup>C242S</sup> | DMEM |
| ASH3 | Human | *PTEN*<sup>F241fs</sup>, *NRAS*<sup>Q61R</sup>, *Tp53*<sup>H179R</sup> | RPMI |
| SW1736 | Human | *BRAF*<sup>V600E</sup>, *TP53*<sup>Q193*</sup> | RPMI |

### Fluorescently labeled cell lines lines

Cells were transduced with lentiviral constructs encoding Luciferase and tdTomato (Addgene #72486) or the cell cycle marker PIP-FUCCI (Addgene #118616). Transduced cells were selected by FACS sorting.

### Cell line preparation and injection

All cell lines used in this study were allowed to expand for 1-2 weeks before harvesting. At 80% of confluence, cells were trypsinized, collected, and resuspended in RPMI1640 without EDTA. A cell suspension at 0.25-0.5 × 10^6^ cells/μL was prepared for injection. Zebrafish larvae xenotransplants injections were performed as previously reported, with some brief alterations (8). At 2dpf, larvae were collected and dechorionated using forceps, anesthetized in 0.016% tricaine (MS-222), and 300-500 cells were injected into the perivitelline space (PVS). Fish were transferred to a plate with E3 to recover for 15-20 minutes before being placed in a 34°C incubator for the remainder of the experiment. At 1dpi the xenotransplant larvae were screened for the presence of cancer cells in the PVS, and larvae without cells or larvae that were improperly injected were discarded.

### I*n-vivo* live imaging

At 1dpi and or 3dpi larvae were anesthetized in 0.016% tricaine and used for non-invasive live imaging at stereomicroscope or spinning disk confocal. All imaging was performed using a zWEDGI device, designed to orient and entrap zebrafish for long-term time-lapse microscopy (6). For long-term imaging, 2% low melting point agarose (Sigma-Aldrich) was placed at loading chamber of the ZWEDGI to retain larvae in the correct orientation. Additional 0.016% tricaine was added as needed over the course of long-term imaging. Images were all acquired with live larvae apart from the EDU assay. Images were acquired on a spinning disk confocal microscope (CSU-W, Yokogawa) with a confocal scanhead on a Zeiss Observer Z.1 inverted microscope equipped with Photometrics Evolve EMCCD camera, and an EC Plan Neofluar NA 0.5/20x air and 0.5/40x air objective, z-stacks, 1.5 μm optical sections. For time-lapse movies, images were taken with a NA 0.5/40x air objective, z-stacks, 1.5 μm optical sections, every 15 minutes for 48 hours.

### Drug treatments

At 1 dpi zebrafish xenotransplants were screened for masses and randomly distributed into treatment groups with 30 larvae per condition, 0.1% DMSO control or 1μM AZD2014 (Vistusertib). Larvae were treated for 3 days in 6-well plates and treatments were refreshed daily (3).

### Histology

After live imaging, larvae were euthanized in 0.4% tricaine and fixed in 2 ml round bottom tubes with 10% formalin. Larvae in 10% formalin were mounted in 2% low melting point agarose and positioned flat on their side. The cassettes were processed by Einstein Histopathology Service, embedded in parafilm, sectioned, and stained with H&E. Slides were digitally scanned by Einstein’s Analytical Imaging Facility on 3D HISTECH P250 High-Capacity Slide Scanner.

### EDU

To confirm cancer cell proliferation EDU staining was performed, as previously reported, larvae were incubated in 10μM 5-ethynyl-2’deoxyuridiine (EdU) solution in E3 for 4 hours with gentle rocking (9). Xenografts were then fixed in 4% PFA overnight at 4°C, washed in 1x PBS, and stored in 100% methanol at -20°C. For the EDU staining, samples were rehydrated in a series of methanol dilutions and permeabilized following the manufacturer instructions on Click-iT EdU imaging kit (Life Technologies). Samples were stored in Vectashield (Vector Laboratories) and then imaged at the Nikon CSU spinning disk confocal microscope. The number of EDU positive cells were automatically counted in the larvae tumor area using IMARIS spots function (9). Spots were defined as particles with 9.5 μm of X/Y and Z diameter (9). To quantify EDU positive cells only in the tumor, a surface was created and then the EDU signal was masked to isolate EDU positive cells in the tumor.

### Mass volume 3D rendering on IMARIS

To quantify tumor volume and angiogenic infiltration, the larval tumor area was imaged as previously described and reconstructed using IMARIS Bitplane Software (Version 9.9.1/10.1) rendering mode. To quantify mass volume/area a 3D surface was generated using the tdTomato fluorescent marker that the cancer cells express. All signal outside of the tumor surface was then masked to isolate endothelial cells within the tumor (9). A second surface was then created for the endothelial cells.

### Automatic quantification of PIP-FUCCI cancer cells on IMARIS

To quantify the total number of cancer cells and the tumor volume, the larval tumor area was imaged as previously described and reconstructed using IMARIS Bitplane Software (Version 9.9.1/10.1) rendering mode and the total number of cancer cells were automatically counted using IMARIS spots function. Spots were defined as particles with 9.5 μm diameter X/Y and Z and to quantify mass volume and area a surface for each region/area was created. For the PIP-FUCCI model the spots and surface function were created in both fluorescent channels. To quantify G2/M cells, that should be fluorescent in both channels, spots were defined as particles with less than 4 μm distance from spots in the opposite fluorescent channel.

### Neutrophil quantification on IMARIS

To quantify neutrophil recruitment to the tumor area and neutrophil infiltration of the tumor, the larval tumor area was imaged as previously described and reconstructed using IMARIS Bitplane Software (Version 9.9.1/10.1) rendering mode and the total number of cancer cells were automatically counted using IMARIS spots function (10). All neutrophils in the FOV (field of view) were counted and neutrophils in the tumor were isolated and counted. Neutrophil density was calculated by normalizing the number of neutrophils per tumor volume.

### Statistical Analysis

All experiments were replicated independently two to three times (N) with multiple samples in each replicate (n). For analysis of mass volume/area, vessel volume/area, and PIP-Fucci cell phases an unpaired Mann-Whitney T-test was performed on GraphPad Prism version 9.3 (10). For mass to vessel volume correlation a Pearson correlation was performed using GraphPad Prism version 9.3. All graphical representations were done on GraphPad Prism version 9.3.

## Authors Contribution

CM, ADC and SDO conceived and designed experiments. CM performed experiments. CM performed data analysis. CM and SdO wrote the manuscript. SDO and ADC critically reviewed and edited the manuscript; All authors reviewed and approved the manuscript.

## Acknowledgments

We want to thank Rita Fior for the helpful discussions and all the support and technical advice given during the development of this work. We thank the Zebrafish core facility, Clint Depaolo and Spartak Kalinin, and the Analytical Imaging Facility (AIF), of Albert Einstein College of Medicine. The AIF is funded by the NCI (P30CA013330).

